# Tissue-specific and repeat length-dependent somatic instability of the X-linked dystonia parkinsonism-associated CCCTCT repeat

**DOI:** 10.1101/2022.01.27.478096

**Authors:** Lindsey N. Campion, Alan Mejia Maza, Rachita Yadav, Ellen B. Penney, Micaela G. Murcar, Kevin Correia, Tammy Gillis, Cara Fernandez-Cerado, M. Salvie Velasco-Andrada, G. Paul Legarda, Niecy G. Ganza-Bautista, J. Benedict B. Lagarde, Patrick J. Acuña, Trisha Multhaupt-Buell, Gabrielle Aldykiewicz, Melanie L. Supnet, Jan K. De Guzman, Criscely Go, Nutan Sharma, Edwin L. Munoz, Mark C. Ang, Cid Czarina E. Diesta, D. Cristopher Bragg, Laurie J. Ozelius, Vanessa C. Wheeler

## Abstract

X-linked dystonia-parkinsonism (XDP) is a progressive adult-onset neurodegenerative disorder caused by insertion of a SINE-VNTR-Alu (SVA) retrotransposon in the *TAF1* gene. The SVA retrotransposon contains a CCCTCT hexameric repeat tract of variable length, whose length is inversely correlated with age at onset. This places XDP in a broader class of repeat expansion diseases, characterized by the instability of their causative repeat mutations. Here, we observe similar inverse correlations between CCCTCT repeat length with age at onset and age at death and no obvious correlation with disease duration. To gain insight into repeat instability in XDP we performed comprehensive quantitative analyses of somatic instability of the XDP CCCTCT repeat in blood and in seventeen brain regions from affected males. Our findings reveal repeat length-dependent and expansion-based instability of the XDP CCCTCT repeat, with greater levels of expansion in brain than in blood. The brain exhibits regional-specific patterns of instability that are broadly similar across individuals, with cerebellum exhibiting low instability and cortical regions exhibiting relatively high instability. The spectrum of somatic instability in the brain includes a high proportion of moderate repeat length changes of up to 5 repeats, as well as expansions of ∼20->100 repeats and contractions of ∼20-40 repeats at lower frequencies. Comparison with *HTT* CAG repeat instability in postmortem Huntington’s disease brains reveals similar brain region-specific profiles, indicating common *trans*-acting factors that contribute to the instability of both repeats. Analyses in XDP brains of expansion of a different SVA-associated CCCTCT located in the *LIPG* gene, and not known to be disease-associated, reveals repeat length-dependent expansion at overall lower levels relative to the XDP CCCTCT repeat, suggesting that expansion propensity may be modified by local chromatin structure. Together, the data support a role for repeat length-dependent somatic expansion in the process(es) driving the onset of XDP and prompt further investigation into repeat dynamics and the relationship to disease.

## Introduction

X-linked dystonia parkinsonism (XDP, OMIM314250) is a progressive and fatal adult-onset neurodegenerative disease endemic to the island of Panay, Philippines [1,2]. The clinical phenotype most commonly described consists of an initial presentation of focal dystonia that spreads to multiple body regions and combines with, or is replaced by, parkinsonism that predominates 10-15 years after onset [1,3,4]. The average age of symptom onset is 39-40 years, though the age at onset (AAO) can differ widely (12 to 64 years) [1,2]. XDP principally affects males, with a frequency of 5.74 cases per 100,000 individuals in Panay, though female carriers are reported to have symptoms in a few cases [1,5]. Limited neuropathological studies of post mortem XDP patient brain tissue have revealed changes to the neostriatum that include selective loss of medium-spiny neurons (MSNs) [6–8] as seen in Huntington’s disease (HD, OMIM 143100) [9]. A handful of neuropathology studies also provides evidence for pathology outside the neostriatum [10,11]. Neuroimaging has demonstrated neostriatal changes, notably atrophy of the caudate and putamen [12–16] as well as changes in cortex, cerebellum, brainstem and globus pallidus [10,12,15].

Genetic linkage and refined mapping localized the causal locus of XDP to the X-chromosome [11,17–19], with recent work characterizing a thirteen-marker haplotype shared by all probands defining a minimal critical region of 219.7 kb with TATA-binding-protein (TBP)-associated factor-1 (*TAF1*) being the only gene within this region [20]. Among the thirteen disease-specific variants is a ∼2.6 kb SINE-VNTR-Alu (SVA)-type retrotransposon [21] inserted in intron 32 of *TAF1* [19]. XDP patient tissues and cell lines exhibit reduced *TAF1* expression [19,20,22–25] as well as aberrant splicing that results in partial retention of intronic sequence proximal to the SVA insertion [20]. Reduced *TAF1* expression, intron retention and aberrant splicing can be rescued by excision of the SVA [20,23], suggesting that SVA-mediated *TAF1* transcriptional dysregulation may contribute to disease pathogenesis. The 5’ end of the SVA contains a hexameric CCCTCT repeat tract that varies in length from 30 to 55 repeats [4,26]. Notably, repeat length is inversely correlated with AAO, as seen in other disorders caused by expanded microsatellite repeats [27], suggesting a critical role of CCCTCT repeat length in XDP pathogenesis. The length of the repeat was also associated with transcriptional activity *in vitro* [4] and its length inversely correlated with *TAF1* expression in patient blood samples [26]. A common characteristic of repeat expansion disorders is the instability of the disease-associated repeat, both in the germline and in somatic tissues, where in the latter the repeat tends to expand in a length-dependent and tissue-specific manner [27–29]. In HD, genetic studies have provided strong evidence that somatic expansion of the *HTT* CAG repeat drives the rate of disease onset [30–32]. Studies of other repeat expansion diseases indicate that somatic expansion is a likely common mechanism driving pathogenesis [28,33–36]. Significantly, a recent genome-wide association study (GWAS) for modifiers of XDP [16] identified genes (*MSH3, PMS2*) with known roles in in repeat instability [31,37–39] that also modify HD [31,40], indicating that a common mechanism at the level of repeat instability extends to XDP. The XDP CCCTCT repeat exhibits intergenerational instability, with repeat length tending to increase in transmissions from mothers and to decrease in transmissions from fathers [4,26]. Patient cell lines show limited repeat instability [4,26], while investigation of a small number of XDP individuals has provided evidence of somatic repeat expansion in post-mortem brain [26,41].

Here, to gain a deeper understanding of somatic instability in XDP we have performed an extensive quantitative characterization of XDP CCCTCT repeat instability in blood, and in up to 17 brain regions from 41 XDP individuals. Our findings reveal repeat length-and tissue-dependent CCCTCT repeat expansion, suggesting that somatic expansion underlies the repeat length-dependent clinical onset of XDP.

## Materials and Methods

### XDP Patients and Sample Collection

#### Blood

Patients recruited for this study included individuals with XDP evaluated at Massachusetts General Hospital (MGH) (Boston, MA, USA), Jose R. Reyes Memorial Medical Center (JRRMMC) (Manila, Philippines), and regional clinics on the island of Panay (Panay, Philippines). All participants provided written informed consent, and the study was approved by local Institutional Review Boards (IRBs) at both MGH and JRRMMC. Patients enrolled were subjected to comprehensive neurological examinations and blood collection [42]. This study also included archival DNA specimens; collection methods and the clinical characterization of donor subjects who provided these specimens have been previously described [18]. Genomic DNA (gDNA) was extracted from blood using the Gentra Puregene kit (Qiagen). Enrolled patients were confirmed to be positive for the XDP mutation by PCR amplification for a known 48 bp deletion haplotype marker as previously described [4,43]. Blood samples from 266 male XDP patients with known AAO were evaluated for correlation with repeat length. Somatic instability was analyzed in 164 blood samples, representing a subset of male XDP patients included in the cohort above.

#### Brain

Post-mortem brain tissue from XDP patients (n=41; 40 with age at onset and death) was obtained in collaboration with the Collaborative Center for XDP (CCXDP), at MGH (Boston, MA, USA), Makati Medical Center (Makati City, Philippines), and the Sunshine Care Foundation (Panay, Philippines). Detailed descriptions of all methods related to donor consent, brain collection and tissue processing have been previously reported [44] and the use of XDP patient post-mortem brain tissue and all study procedures were approved by Institutional Review Boards at Makati Medical Center (Makati City, Philippines) and MGH (Boston, MA, USA). Genomic DNA was extracted from different brain regions using the DNeasy Blood and Tissue Kit (Qiagen), according to manufacturer’s instructions and with the following modifications: samples were incubated in buffer ATL and Proteinase K overnight at 56°C; washes AW1 and AW2 were repeated; DNA was eluted in 100 μl of Qiagen Elution Buffer, preheated to 56°C, applied to the spin columns, and incubated at room temperature for 10 minutes before centrifugation. The sample was run through the spin column a second time before final centrifugation. The presence of the XDP mutation in each brain was confirmed as above.

### Determination of XDP and *LIPG* CCCTCT repeat lengths and expansion indices

To determine the length of XDP and *LIPG* SVA CCCTCT repeats in blood and postmortem brain regions, we used fluorescent PCR-based assays, with the primers and conditions outlined in Additional File 1: Table S1. Both protocols used 125ng of gDNA per reaction, in a 25 µl reaction volume with buffer and dNTPs provided with the PrimeSTAR GXL polymerase (Takara) according to the manufacturer’s protocol, and as previously described for the XDP repeat [4]. Following PCR, aliquots of each product were resolved via electrophoresis in agarose gels to confirm amplification of the SVA repeat sequence and then run on the ABI 3730 DNA sequencer (Applied Biosystems) with GeneScan 500 LIZ as internal size standard, and the data analyzed using GeneMapper v5 (Applied Biosystems) [4]. Repeat amplification resulted in a distribution of fragments separated by 6 bp and repeat size was defined as the tallest peak in this distribution. XDP repeat size was assigned relative to a sequenced control and *LIPG* repeat size calculated based on fragment length (bp). To quantify XDP and *LIPG* CCCTCT repeat expansion, we generated an expansion index from the GeneMapper peak height data as previously described [45], using a 5% relative peak height threshold cut-off (*i*.*e*. excluding peaks whose height is less than 5% of the height of the modal allele). Because *LIPG* is autosomal, most individuals had two distinguishable allele lengths. In many cases, alleles were sufficiently separated to allow quantification of expansion peaks from each. In some individuals, when the alleles were too close, we only captured the expansion index from one allele.

### Small pool-PCR Southern blot analyses

1 µg of gDNA was digested with HaeIII (37°C for 12 hours) and the enzyme subsequently inactivated at 80°C for 20 minutes. Serial dilutions were made in water to a final concentration of 90 pg/µl and 1µl (approx. 30 genome equivalents, g.e.) was used for PCR amplification using a non-FAM-labeled version of the XDP SVA hexamer primers with the small pool-PCR conditions outlined in Additional File 1: Table S1. For each sample, PCR amplifications of 36 replicates of 90 pg gDNA and 8 DNA-negative PCR controls were carried out in a 25 µl reaction volume with buffer and dNTPs provided with the PrimeSTAR GXL polymerase (Takara) according to the manufacturer’s protocol. 10 µl of each PCR product were run in 2% agarose gels alongside digoxigenin (DIG)-labeled size markers VII and VIII (Roche), for 16 hours at 50 V then transferred to a positively charged nylon membrane (Roche) by common squash-blotting technique [26,46]. The membrane was hybridized with 5 pmol/ml of a 5’ DIG-labeled (AGAGGG)10 probe (Sigma) in DIG Easy Hybridization Solution (Sigma) overnight at 45°C and then washed twice each with 2 X SSC, 0.1 % SDS at room temperature for 5 minutes, 0.1 X SSC, 0.1% SDS at 68°C for 20 minutes, and 0.1 X SSC, 0.5% SDS at 68°C for 20 minutes. DIG detection was carried out using the DIG Luminescent Detection system (Sigma) with CPSD substrate according to the manufacturer’s instructions.

### Single molecule small pool-PCR sizing

1 µg gDNA was digested with HaeIII as above and the DNA serially diluted to a range of concentrations spanning 3 pg/ul to 300 pg/ul corresponding to approximately 0.5-100 diploid g.e/µl, respectively. For each sample, at least 10 PCR reactions of 1 µl DNA inputs were run for each dilution and resolved using the ABI 3730 DNA Sequencer. Poisson analysis was used to determine empirically for each sample the concentration of DNA that resulted in single molecule PCR amplification, *i*.*e*. the concentration that resulted in ∼33% of all DNA input reactions having no product. We then ran, for each sample, at least three 96-well plates, each consisting of 72 replicates of the optimized single molecule amplifiable DNA amount, 18 DNA-negative PCR controls, 5 XDP repeat sizing controls, and one empty well for machine control. PCR conditions for small pool-PCR were as described in Additional File 1: Table S1, and CCCTCT repeat size was determined as described above. Allele lengths between 330 bp and 560 bp (about 32-70 repeats) could be accurately determined based on the known repeat sizing controls. For PCR products falling outside of this range we estimated repeat length based on molecular weight. All peaks with heights >=150 were sized, and for each plate we verified that all of the no-DNA input wells were negative and that at least 1/3 of the DNA input wells were negative.

### HD sample data

In this study we used *HTT* CAG repeat expansion data previously generated and reported from 8 postmortem brain regions from three HD individuals (HD1-3; CAG repeats 43/16, 44/17, 53/19) obtained from the New York Brain Bank under an approved MGH IRB protocol [29]. The data from a subset of eight tissues used in this study were chosen because they were identical or as close as possible to the XDP brain regions from our XDP cohort. Regions compared were: BA9 (HD and XDP), BA17 (HD) and occipital cortex (XDP), caudate, accumbens and putamen (HD) and caudate (XDP), cerebellum (HD and XDP), cingulate gyrus (HD and XDP), globus pallidus putamen (HD) and putamen (XDP), hippocampal formation (HD) and hippocampus (XDP), subthalamic nucleus (HD and XDP), temporal pole (HD and XDP). For simplicity, in Figure 5c we refer to all the regions according to the XDP labels. Somatic *HTT* CAG expansion indices were determined for this study using a 5% relative peak height threshold cut-off for comparison to the 5% threshold XDP CCCTCT expansion indices.

### Statistical analysis

Data analysis and plots were generated using R/RStudio V.1.3. (https://cran.r-project.org/mirrors.html). Linear regression, stacked bars and scatter plots were generated using ggplot2 package (https://www.rdocumentation.org/packages/ggplot2/versions/3.3.5). Pearson or Spearman coefficients were determined using ggscatter package and used as appropriate when data distribution was Normal or not, respectively. The heatmap was generated from a scaled dataset using the heatmaply package followed by a clusterization method, based on Manhattan distance https://cran.r-project.org/web/packages/heatmaply/vignettes/heatmaply.html. Multiple pairwise comparison test was performed using Wilcoxon rank-sum test followed by Bonferroni Post Hoc method for *P*-value adjustment. X^2^ test was used to compare the numbers of events from single molecule SP-PCR data across brain tissues. *P*-value < 0.05 was considered significant.

## Results

### XDP CCCTCT repeat length inversely correlates with ages at onset and death but not with disease duration

We previously demonstrated in a cohort of 140 XDP males that CCCTCT repeat length in blood was inversely correlated with AAO [4]. This observation was subsequently confirmed in an independent cohort of 295 individuals [26]. Here, we have used an expanded dataset from our original sample of 140 comprising blood (n=266) and brain (n=40) DNA samples from clinically confirmed male XDP patients to examine further the relationship between CCCTCT repeat length and AAO, as well as age at death (AAD) and disease duration, defined as AAD minus AAO (Fig. 1). In these analyses, brain repeat length was determined in 40 postmortem samples with AAO (n=39 in cerebellum and n=1 in occipital cortex where cerebellum was not available). Both blood and brain tissue were available for 21 individuals; of these, blood and brain (cerebellar) repeat lengths were identical in 17 individuals and differed by one repeat in 4 individuals (17-012, 19-017, 19-021, 21-031; Additional file 1: Table S2). Mean (±SD) repeat lengths in blood and brain were 41.6 ± 3.9 (range:34-53) and 41.8 ± 4.6 (range:34-55), respectively. Mean (±SD) AAO of the blood and brain samples were 41.4 ± 8.3 (range:18-65) and 41.4 ± 8.7 (range:26-59) years, respectively. Blood repeat length inversely correlated with AAO and explained ∼45% of the AAO variability (*P=*7.7*e*-36; Fig. 1a, red dots), consisted with previous studies [4,16,26]. A similar correlation was observed between brain repeat length and AAO, with repeat length explaining ∼55% of the AAO variability (*P*=4.7*e*-08; Fig. 1a, blue dots). There was no difference in AAO-repeat length correlation between individuals exhibiting primarily dystonia at onset (N=194) and those exhibiting primarily parkinsonism at onset (N=43) (Additional file 2: Fig.S1), consistent with previous observations [26]. Both AAO and AAD was available for 68 individuals, 28 of whom had blood repeat sizing and 40 of whom had brain repeat sizing. As repeat length was largely identical between brain and blood for the individuals with both measures, we used a combined blood and brain dataset from these 68 individuals to examine relationships between repeat length and AAO, AAD or duration (Fig1. b-d). Mean (±SD) repeat length in these 68 individuals was 41.6 ± 4.4 (range:34-55), mean (±SD) AAO was 41.7 ± 4.4 (range:26-64), and mean (±SD) AAD was 50.7 ± 9.5 (range:30-69) years. Repeat length inversely correlated with AAO and AAD, explaining ∼53% (*P*=2.3*e*-12) and ∼42% (*P*=2.5*e*-09) of the AAO and AAD variability, respectively (Fig. 1b-c). In contrast, we found no significant correlation between repeat length and disease duration (AAD-AAO) (Fig. 1d). These data indicate that the length of the CCCTCT repeat is critical for process(es) driving XDP onset and death that ensues, though has no obvious effect or a weaker effect on duration.

**Fig. 1.**
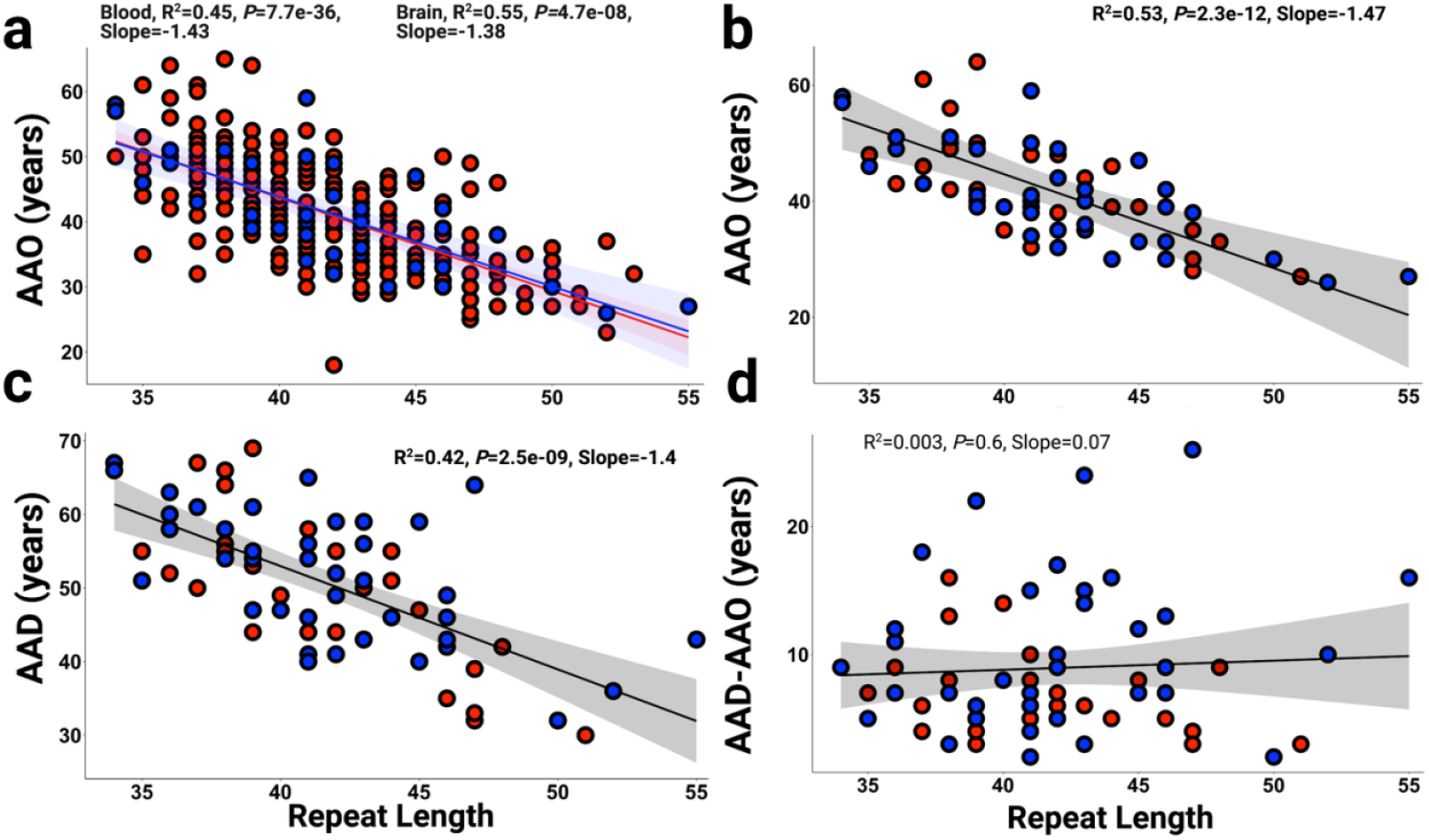
Length of the CCCTCT repeat correlates with AAO and AAD in male XDP patients. **(a)** Inverse correlations between CCCTCT repeat length in blood (red dots and line, n=266) and brain (blue dots and line, n=40) with AAO. **b** Inverse correlation between CCCTCT repeat length determined in a subgroup of blood and brain samples from deceased XDP patients (blood n=28; brain n=40) and AAO. **c** Inverse correlation between CCCTCT repeat length determined in a subgroup of blood and brain samples from deceased XDP patients (blood n=28; brain n=40) and AAD. **d** Length of the CCCTCT repeat determined in the subgroup of blood and brain samples from deceased XDP patients (blood n=28; brain n=40) is not correlated with disease duration (AAD-AAO, n=68). AAO, age at onset; AAD, age at death. Brain repeat lengths were determined in cerebellum (n=39) or occipital cortex (n=1). In a-d, blood (red dots) and brain samples (blue dots).

### The XDP CCCTCT repeat exhibits tissue-and repeat length-dependent somatic expansion

The variation in repeat length between individuals reflects the instability of the CCCTCT repeat in germline transmissions [4,26]. To gain insight into CCCTCT repeat instability in somatic tissues we have examined repeat length variation in blood (n=164) and postmortem brain (n=41) from affected males. In the brain, we analyzed between 1 and 17 brain regions in 41 individuals, including cerebellum only in 17 individuals and occipital cortex only in one individual (Additional File 1: Table S3). The XDP CCCTCT repeat was PCR-amplified using a previously established genotyping assay for repeat sizing [4]. PCR amplification of the repeat results in a distribution of fragment sizes, with repeat length determined as the modal allele in the distribution. Of the 23 postmortem samples in which multiple brain regions were analyzed, 4 (17-012, 17-17, 19-017 and 21-031) exhibited variation by one repeat unit (Additional File 2: Fig. S2) while in 19 individuals the modal repeat length was identical in all brain regions analyzed. Therefore, XDP CCCTCT repeat instability is not substantially reflected in differences in modal repeat length of the repeat-containing PCR amplicons.

We then analyzed XDP CCCTCT instability by quantifying an expansion index from repeat length distributions of GeneMapper outputs of the repeat-containing PCR products [45]. This relatively high throughput method is sensitive to subtle differences in repeat instability that are captured in the majority of alleles. Examples of GeneMapper traces from different tissues are shown in Additional File 2: Fig. S3. The peaks to the left of the modal allele are largely due to PCR slippage, and therefore we quantified only the expansion peaks to the right of the modal allele. These peaks are variable between tissues and are the result of somatic repeat length variation. Expansion indices in blood and brain regions are shown in Fig. 2a, ordered from left to right by the median expansion index per tissue. Very low levels of XDP CCCTCT expansion were detected in blood (median expansion index = 0.19, interquartile range [IQR]= 0.22). In contrast, all brain regions exhibited expansion indices that were significantly greater than those in blood (*P*<0.05: Wilcoxon rank-sum tests with Bonferroni correction; Additional File 1: Table S4). Of the brain regions analyzed, cerebellum had the lowest expansion index (median expansion index = 0.77, interquartile range [IQR]= 0.32), while occipital cortex exhibited the highest expansion index (median expansion index = 1.59, interquartile range [IQR]= 0.7). Replicate PCR amplifications from the same DNA samples demonstrated that differences between brain regions are not due to technical variation (Additional File 2: Fig. S4). Statistically significant differences in expansion indices (*P*<0.05: Wilcoxon rank sum tests with Bonferroni correction) were observed between some of the brain regions, most notably in comparisons with cerebellum or occipital cortex (Additional File 1: Table S4). Overall, there appeared to be a tendency towards higher expansion indices in cortical regions (cingulate gyrus, prefrontal cortex (BA9), parietal cortex, insula, temporal pole and occipital cortex) than subcortical areas (cerebellum, caudate, substantia nigra, inferior olivary nucleus, red nucleus, medial thalamus, hippocampus, putamen, lateral thalamus, deep cerebellar nuclei, sub-thalamic nucleus). Of the subcortical structures, there was no obvious distinction in expansion indices between forebrain (caudate, putamen, hippocampus, thalamus, subthalamic nucleus), midbrain (red nucleus) or hindbrain (deep cerebellar nuclei, inferior olivary nucleus) regions, with the exception of cerebellum (Fig. 2a). Due to the considerable variation in repeat expansion between individuals, we further evaluated tissue patterns of expansion by performing hierarchical clustering on a heatmap plot based on scaled expansion index values (Fig. 2b). The heatmap revealed similar patterns of brain region-specific expansion across individuals and distinguished two major clusters comprised of cortical and subcortical brain areas (Fig. 2b).

**Fig. 2.**
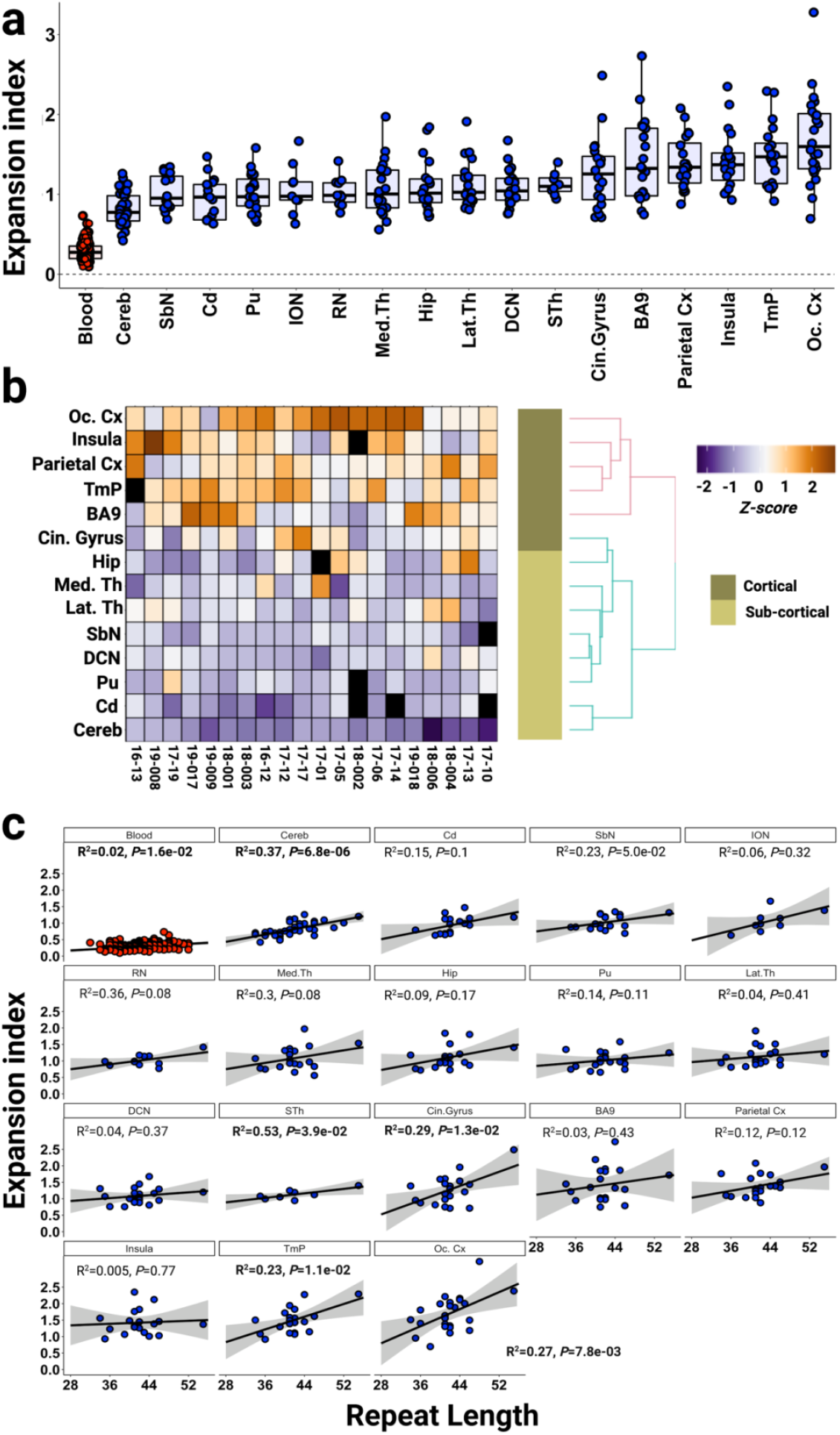
XDP CCCTCT repeat expansion index in blood and brain regions. **a** Distribution of expansion indices ranked by median values in blood and brain regions. Box-whisker plots show median ± interquartile range (IQR) and dots show values in individual patient samples. **b** Heatmap of expansion indices values in different individuals (rows), scaled (z-score) across brain tissues (columns). To avoid poor normalization during scaling, brain regions with fewer than 12 measures and individuals with fewer than 6 tissue samples were excluded (Additional File 2: Fig. S4). Brain regions with no measurement are represented as black boxes. **c** Linear regression analyses showing relationships between CCCTCT repeat length and expansion index in each tissue. The regression equations shown in bold font highlight those tissues (blood, cerebellum, subthalamic nuclei, cingulate gyrus, temporal pole, occipital cortex) showing a significant association of expansion index with repeat length. Grey shaded areas show 95% confidence interval. Blood (n=164), Cereb=cerebellum (n=40), Cd=caudate (n=17), SbN=substantia nigra (n=19), ION = inferior olivary nucleus (n=9), RN = red nucleus (n=11), Med.Th = medial thalamus (n=20), Hip = hippocampus (n=19), Pu = putamen (n=19), Lat.Th = lateral thalamus (n=20), DCN = deep cerebellar nuclei (n=21), STh = subthalamic nucleus (n=8), Cin.Gyrus = cingulate gyrus (n=20), BA9 = frontal cortex Brodmann area 9 (n=21), Parietal Cx = parietal cortex (n=20), Insula = insular cortex (n=19), TmP = temporal pole (n=20), Oc. Cx = occipital cortex (n=24).

As individuals differ in their repeat length, we investigated the extent to which repeat length might explain the variation in expansion index within any one tissue (Fig. 2c). Overall, the data showed positive correlations between expansion index and repeat length that were statistically significant in a subset of the tissues (blood, cerebellum, subthalamic nuclei, cingulate gyrus, temporal pole, occipital cortex). The proportion of the variation in expansion index explained by repeat length varied from 2% in blood to 45% in the red nucleus. The cerebellum exhibited the most significant correlation (*P*=6.8×10^−6^), with repeat length explaining 37% of the expansion index variation. The various strengths of the associations with repeat length likely differ as a function of sample number, the magnitude of the instability, and the cell type heterogeneity in each tissue piece that is sampled. *e*.*g*. blood shows minimal repeat expansion, limiting the sensitivity to detect biological variation. In cerebellum, the relatively strong association with repeat length is likely contributed by both cell type homogeneity - 99% of all cerebellar neurons are granule cells - and the greater number of cerebellar samples relative to the other brain regions.

Together, these data demonstrate greater somatic expansion of the XDP CCCTCT repeat in the brain than in blood as well as brain region-specific propensities for expansion that are similar across individuals. Significantly, we show that somatic CCCTCT expansion is dependent on repeat length, consistent with a contribution of somatic expansion to the onset of disease.

### The XDP CCCTCT repeat exhibits large repeat length changes and expansion-biased instability in the brain

Analysis of repeat instability in fragment sizing data obtained from PCR-amplified “bulk” genomic DNA, as above, is limited by the lack of sensitivity to detect rare alleles and an upper limit for accurate fragment sizing of ∼330-560 base pairs, equating to ∼ 32-70 CCCTCT repeats. Further, while allele length distributions can be quantified in the PCR products, as with the expansion index metric, this may not accurately reflect the distribution of allele lengths present in genomic DNA due to contraction bias inherent to the PCR. Therefore, to investigate more fully the spectrum of repeat length mosaicism in XDP brains we employed two small pool-PCR (SP-PCR) approaches, providing the sensitivity to detect rare somatic events and to quantify allele size distributions. We analyzed a subset of the brain tissues, sampling across regions (occipital cortex, caudate, putamen, cerebellum) exhibiting a range of instabilities as determined from the GeneMapper-based analysis above, and across individuals with a range of repeat lengths (17-17: 54/55 repeats, 19-008: 41 repeats, 18-006: 35 repeats; Fig. 3, Table 1).

**Table 1.** Summary of XDP CCCTCT repeat sizing and instability analyses.

| Sample | Repeat<br>length<br>(standard<br>genotyping) | Expansion<br>index | Small pool-PCR<br>Southern blot<br>Highest/lowest repeat<br>lengths | Single molecule input small pool-PCR |  |  |  |
| --- | --- | --- | --- | --- | --- | --- | --- |
|  |  |  |  | Number<br>of alleles<br>sampled | Mean<br>repeat<br>length | Modal<br>repeat<br>length | Highest/lowest<br>repeat<br>lengths |
| 17-17 Cerebellum | 55 | 1.205 | 82/31 | 121 | 60 | 56 | 101/25 |
| 17-17 Occipital Cortex | 54 | 2.383 | 129/36 | 211 | 55 | 54 | 100/24 |
| 17-17 Putamen | 54 | 1.247 | 128/30 | 147 | 55 | 54 | 71/50 |
| 17-17 Caudate | 54 | 1.1809 | 97/42 | 243 | 54 | 54 | 84/19 |
| 18-006 Occipital Cortex | 35 | 0.9524 | 149/22 | 182 | 37 | 35 | 60/26 |
| 19-008 Occipital Cortex | 41 | 1.305 | 77/31 | 168 | 42 | 41 | 53/21 |
| 19-008 Cerebellum | 41 | 1.111 |  | 141 | 42 | 41 | 50/26 |
| 19-008 Caudate | 41 | 1.124 |  | 142 | 42 | 41 | 49/37 |
| 19-008 Putamen | 41 | 1.061 |  | 164 | 42 | 41 | 47/35 |

**Fig. 3.**
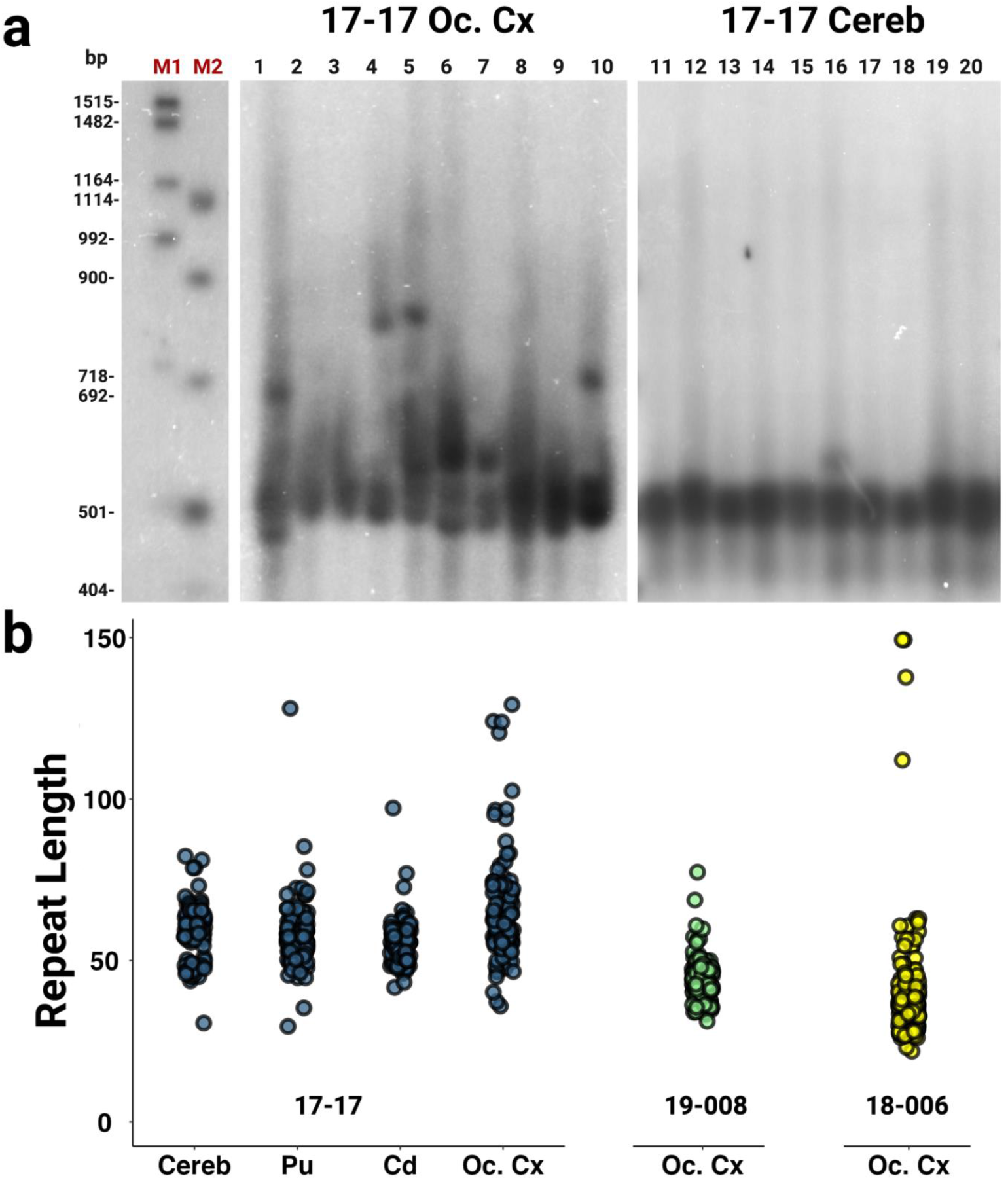
Southern blot images and estimated repeat lengths. **a** Representative Southern blot images for 17-17 occipital cortex (Oc. Cx) (lanes 1-10) and cerebellum (Cereb) (lanes 11-20) illustrating the varying degree of instability across brain regions. Each lane represents PCR amplification of ∼30 g.e. M1 and M2 size markers are DIG VII and VIII, respectively and are shown with the corresponding base pair lengths. **b** Estimated CCCTCT repeat lengths based on distance migrated relative to the M1 and M2 markers. Repeat size data for each sample are obtained from 36 replicates (individual lanes), each with ∼30 g.e. input DNA amount.

We first performed SP-PCR in conjunction with Southern blot detection, diluting the genomic DNA to approximately 30 genome equivalents (g.e) prior to PCR amplification of the CCCTCT repeat. Examples of the Southern blots are shown in Fig. 3a and a summary of the data is provided in Fig. 3b and Table 1, the latter indicating the approximate highest and lowest repeat lengths detectable for each sample. The approximate length ranges for the greatest density of signal on the Southern blots encompassed the repeat sizes determined by standard genotyping (Fig. 3b, Table 1). Notably, all samples showed distinct additional bands reflecting expansions or contractions, with a bias towards expansions. The largest alleles detected across all samples ranged from ∼77 to ∼149 repeats with increases in length relative to those determined by standard genotyping ranging from ∼27 to >100 units (Table 1). The smallest alleles detected ranged from ∼22 to ∼42 repeats, representing ∼13-24 unit decreases relative to genotyped repeat lengths. The highest and lowest approximate repeat lengths detected were found in 18-006 occipital cortex (149 and 22, respectively) despite this sample having the shortest genotyped repeat and smallest expansion index (Table 1). Among the different tissues from individual 17-17 (54 repeats), occipital cortex exhibited the most instability, with repeats ranging from 36 to 129. Cerebellum was the most stable of these tissues, but nevertheless did show evidence for alleles ranging from 31 to as high as 82 repeats. Caudate and putamen exhibited degrees of mosaicism between those of occipital cortex and cerebellum. In occipital cortex from 19-008 (41 repeats) we detected a range of repeat lengths from 31-77. In general, qualitative patterns of instability observed on the Southern blots approximately parallel quantitative differences in expansion indices (Table 1, Fig. 3) but highlight the occurrence of rarer somatic events that are not detected in the bulk PCR-based analyses.

While the SP-PCR Southern blot analyses allow detection of large repeat length changes, input DNA amounts of multiple genomes do not allow for quantitative analyses of repeat length distributions in these samples as signals from individual amplification products are not necessarily distinguishable. To quantify repeat length distributions, we therefore performed SP-PCR of single input molecules. We targeted ∼120-240 individual molecules per sample (Table 1) with the aim of capturing somatic events that occurred at a frequency of ∼0.5-1%, and sized individual PCR products on the ABI sequencer to achieve single repeat resolution. It should be noted that fragment sizing of SP-PCR products has the same sizing limitations as bulk PCR and thus we were not able to assess the very large rare expansions that were seen on Southern blots. We examined the same brain samples as for the Southern blot-based analyses and extended the single molecule analyses to include putamen, caudate, cerebellum in addition to occipital cortex from 19-008 (Table 1, Fig.4 and Additional File 1: Table S5). These data revealed a high proportion of alleles with lengths either expanded or contracted relative to the modal repeat length (Fig.4b). Note that the modal repeat length in the single molecule input SP-PCR data was identical to the repeat length determined by standard genotyping of bulk genomic DNA with the exception of 17-17 cerebellum where SP-PCR modal allele was greater by one repeat (Table 1). Across these samples 65% to 84% (mean 74%) of alleles deviated from the modal allele length. The frequency of expansions ranged from 30% to 58% (mean 49%) while the frequency of contractions was lower overall, ranging from 16% to 45% (mean 26%) (Fig.4a, Table S5). The relative frequencies of contracted, modal and expanded alleles differed across the four brain regions of individual 17-17 (Chi^2^=33.30, df=6, *P*<0.0001) with a relatively high proportion of expansions in occipital cortex and a relatively low proportion of expansions in cerebellum. Relative frequencies of contracted, modal and expanded alleles were not significantly different between the four brain regions of individual 19-008 (Chi^2^=8.882, df=6, *P*=0.1803) but differed significantly between occipital cortices of the three individuals (Chi^2^=12.52, df=4, *P*=0.0139). The majority of the expanded alleles were 1-4 repeat units, with expansions of 5 or more repeats occurring in 2%-18% of alleles (mean 9%) and expansions of 20 or more repeats occurring in 0%-12% of alleles (mean 2%) (Additional File 1: Table S5). The majority of the contracted alleles were also in the range of 1-4 units, with contractions of 5 or more repeats in 0-11% of alleles (mean 3%) and contractions of 20 or more repeats in 0-3% of alleles (mean 0.8%) (Table S5). Overall, the allele size distributions in the single molecule data capture both the tissue-specific and individual-specific differences in instability that are similarly reflected in the expansion index measure and SP-PCR Southern blot analyses.

**Fig. 4.**
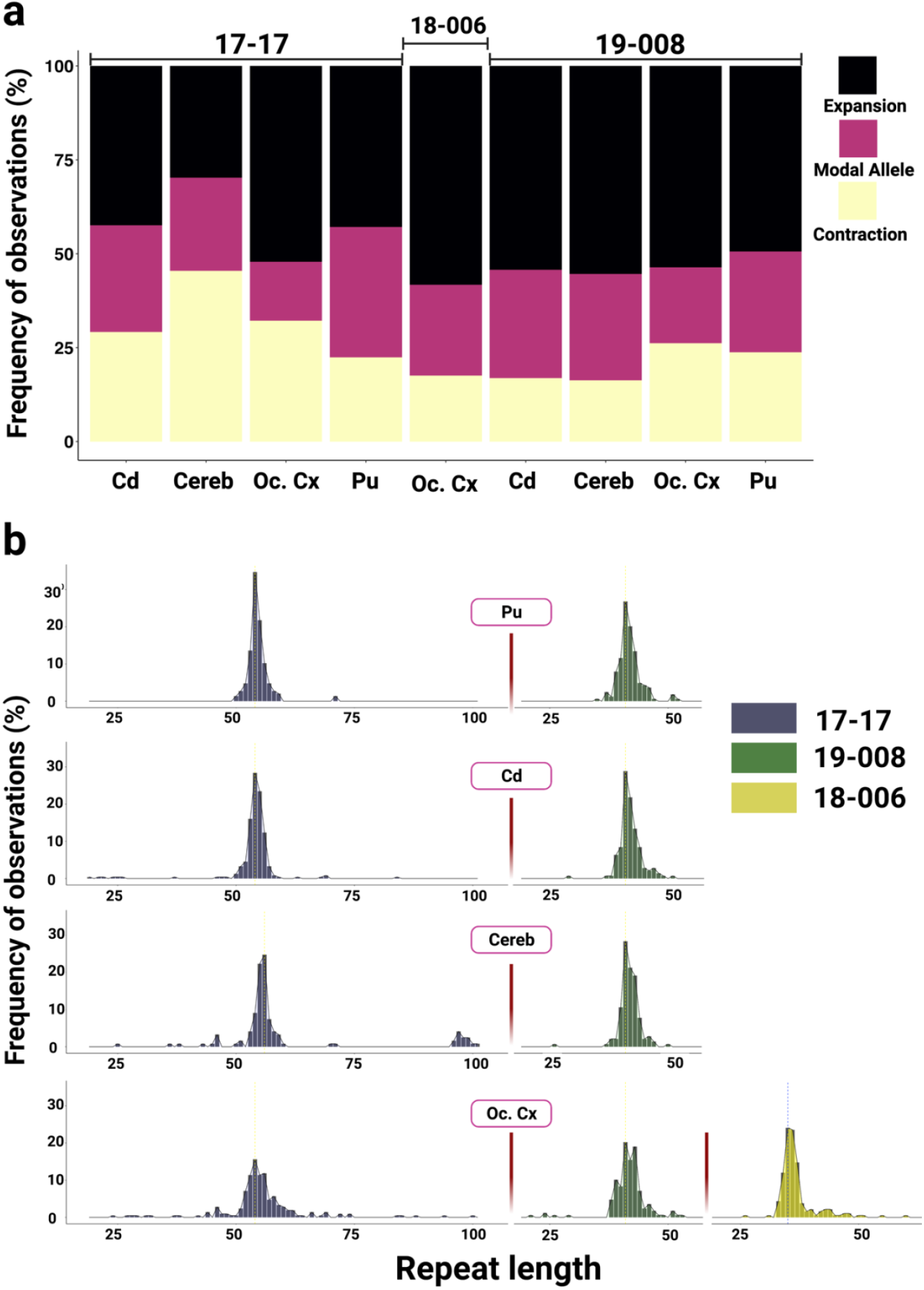
XDP CCCTCT repeat length distributions in brain regions. Repeat lengths were determined by fragment sizing of amplicons obtained in single molecule input PCRs in four tissues (Pu=putamen, Cd=caudate, Cereb= cerebellum, Oc. Cx =occipital cortex) across three patients. **a** Percentages of expansions and contractions compared to the modal allele. **b** Histograms of repeat length frequencies. Data in **a** and **b** were derived from 121-243 single amplifiable molecules for each sample. Refer to Table 1 for summary data derived from these analyses.

### Features of XDP CCCTCT somatic expansion are shared among other microsatellite repeats

To gain additional insight into XDP CCCTCT repeat dynamics we were interested in exploring overlaps with other microsatellite repeats, in particular: 1) a different CCCTCT repeat, and 2) the unstable expanded *HTT* CAG repeat due to shared genetic and pathological features of HD and XDP. No other disease-causing CCCTCT repeats have been described to date, however CCCTCT repeats are common elements of SVA retrotransposons in the human genome [21]. To identify another CCCTCT repeat to study in comparison to the XDP repeat, we defined inclusion criteria as: 1) the repeat is similar in length to the XDP repeat (∼35-50) and 2) the repeat-containing SVA is located in an intron and inserted in reverse orientation relative to the gene transcript, as it is for the XDP SVA. We thus identified a CCCTCT repeat of 39 units in the reference genome (hg19 chr18:47105372-47105605) within an SVA inserted in reverse orientation in intron 5 of the endothelial lipase G gene (*LIPG*), hereafter referred to as the *LIPG* CCCTCT repeat. We first PCR-amplified the *LIPG* CCCTCT repeat from a subset of XDP patient cerebellar DNAs. Repeat length varied from 39 to 71 (median=53, IQR=10), with two repeat lengths distinguishable in some individuals and only one in others (Table S6). We then identified six individuals for analyses of *LIPG* CCCTCT repeat instability across brain regions (cerebellum, caudate, hippocampus, BA9, temporal pole and occipital cortex) that exhibited a range of XDP CCCTCT expansion levels. The six individuals were selected based both on tissue availability and having two *LIPG* CCCTCT repeat lengths sufficiently well-separated to allow quantification of an expansion index from each allele (Additional File 1: Table S6). Examples of GeneMapper outputs of *LIPG* CCCTCT repeat-containing PCR products are shown in Additional File 2: Fig.S5. Quantification of an expansion index across all the brain samples (Fig. 5a) revealed the lowest expansion index in cerebellum (median=0.12, IQR=0.06), and the highest expansion index in caudate (median=0.95, IQR= 0.5), with significantly lower cerebellar expansion indices relative to other brain regions (*P*<0.05: Wilcoxon rank-sum tests with Bonferroni correction, Additional File 1: Table S7). A comparison of *LIPG* and XDP CCCTCT expansion indices (Fig.5a) revealed significantly lower values for cerebellum, hippocampus, BA9, temporal pole and occipital cortex brain regions despite the *LIPG* having longer repeats on average than the XDP repeat (*P*<0.05: Wilcoxon rank-sum test, Additional File 1: Table S7). *LIPG* CCCTCT expansion indices also positively correlated with repeat length with the proportion of the variation in expansion index explained by repeat length varying from 54% in caudate to 69% in the temporal pole (Fig. 5b). It is worth noting that the variability in expansion index as a function of repeat length (R^2^) may be overestimated in these data due to the inclusion of two alleles from the same individual. Overall, despite the small sample size and lower absolute levels of expansion of the *LIPG* repeat compared to the XDP repeat, these data reveal that both repeats share properties of length-dependent expansion being relatively low in cerebellum.

We previously reported, using similar quantitative analyses, tissue-specific patterns of somatic expansion of the *HTT* CAG repeat in HD postmortem brains [29]. To compare tissue-specific instability of the XDP CCCTCT and *HTT* CAG repeats, we plotted mean expansion indices across all patient samples for nine brain regions (BA9, cerebellum, hippocampal formation, temporal pole, putamen, occipital cortex, subthalamic nuclei and caudate) that were shared across the HD study and this XDP study (see Materials and Methods). We found that XDP and *HTT* repeat expansion indices in XDP and HD patient brain tissues, respectively, were highly correlated (correlation coefficient *r*=0.65, *P*=0.0057, Fig. 5c), indicating shared tissue-specific expansion propensities of these two different disease-associated repeats. In contrast, and as indicated in Fig.5a, the XDP and *LIPG* expansion indices are not correlated across the six tissues analyzed (correlation coefficient *r*=0.14, *P*=0.79).

**Fig. 5.**
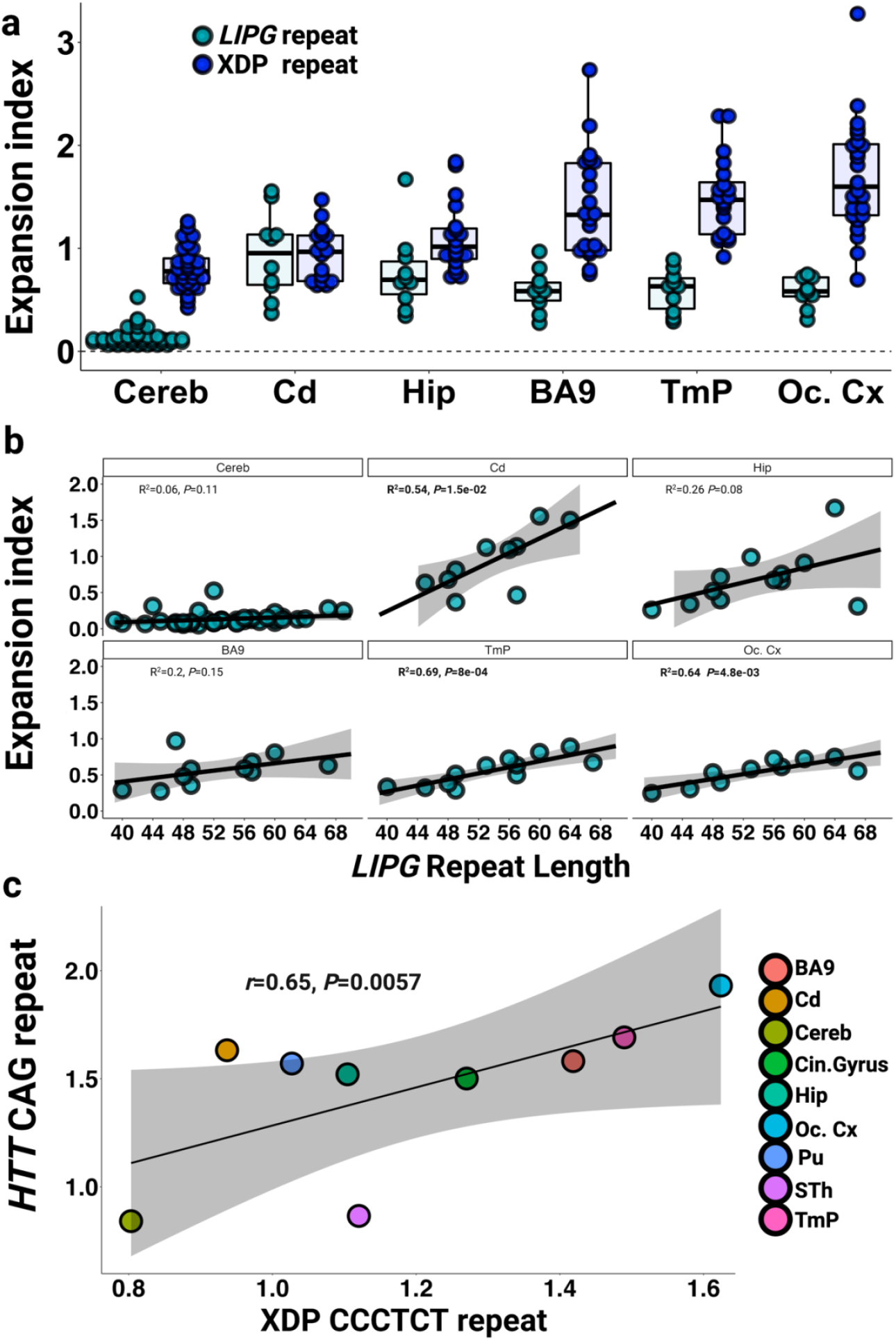
Expansion of the CCCTCT *LIPG* repeat and *HTT* CAG repeat in comparison to the XDP CCCTCT repeat. **a** Distribution of expansion indices of *LIPG* and XDP CCCTCT repeats in XDP postmortem brain tissues. Box-whisker plots show median ± IQR and dots show values for individual alleles. XDP repeat: data are the same as in Fig.2. Refer to Additional File1: Table S3 for sample numbers for each brain region. *LIPG* repeat: Cereb=Cerebellum (n=23 individuals, 40 alleles), Cd=Caudate (n=5 patients, 10 alleles), Hip = Hippocampus (n= 6 patients, 10 alleles), BA9 = frontal cortex Brodmann area 9 (n= 6 patients, 11 alleles), TmP = Temporal pole (n=6 patients, 12 alleles), Oc. Cx = Occipital cortex (n=5 patients, 10 alleles). Note that some alleles that failed QC were excluded. **b** Linear regression analyses showing relationships between *LIPG* CCCCTC repeat length and expansion index in each brain region. The regression equations shown in bold font highlight those tissues (caudate, BA9, temporal pole and occipital cortex) showing a significant association of expansion index with repeat length. Grey shaded areas show 95% confidence interval. **c** Correlation of mean *HTT* CAG expansion index in three HD individuals (Materials and Methods) and mean XDP CCCTCT expansion indices. Refer to Additional File1: Table S3 for sample numbers for each brain region for XDP.

## Discussion

Previous studies have shown that the length of the XDP-associated CCCTCT repeat in blood is inversely correlated with AAO, accounting for ∼50% of the AAO variance [4,26]. There is also evidence for correlations between repeat length and other clinical disease measures [26]. The present study supports and extends these data; in our expanded blood dataset (N=266), repeat length accounted for ∼46% of the variance in AAO, and in as few as 40 individuals we detected a significant correlation between AAO and repeat length measured in brain DNA (R^2^=0.55). The different R^2^ values between the various studies and our cohorts [26,47] may in part be explained by differences in the accuracy in determining AAO and warrants additional investigation. In addition, in a subset of individuals with known AAO and AAD we show for the first time that repeat length is inversely correlated with AAD, with a relationship paralleling that between repeat length and AAO. In contrast, we observed no significant correlation between repeat length and disease duration (the time between onset and death), an observation previously reported in HD [48]. However, this does not preclude a possible stronger effect of repeat length on duration that is counterbalanced by an effect of AAO [49] (*i*.*e*. longer repeat length resulting in a shorter duration, counterbalanced by longer repeat length resulting in earlier AAO and subsequent longer duration). As AAD was only available for 68 individuals in this study, additional patient data will be needed for further dissection of repeat length-dependent relationships with disease duration. Importantly, our data underscore the importance of CCCTCT repeat length in driving the rate of XDP, motivating the investigation of the instability of this repeat tract in somatic cells in patients.

To gain insight into the somatic instability of the XDP-associated CCCTCT repeat we have used multiple methodologies, including single molecule-based analyses, to probe the spectrum of repeat length mosaicism in blood and across seventeen brain regions from XDP patients. We demonstrate that the XDP CCCTCT repeat exhibits extensive somatic mosaicism, notably length-dependent and tissue-specific expansion that is measurable in the bulk of alleles, and the presence of rarer alleles in the brain that can be either substantially contracted or expanded on the order of ∼10s >100 repeats relative to the repeat length determined using standard genotyping. Given the inverse correlation of CCCTCT repeat length with AAO these observations implicate somatic expansion as a driver of the rate of onset of XDP. Notably, a GWAS identified two genes, *MSH3* and *PMS2* as modifiers of the age of onset of XDP [16]. These genes are also modifiers of HD age at onset [31] and encode DNA mismatch repair proteins that modulate the somatic instability of disease-associated trinucleotide repeats, including the *HTT* CAG repeat [31,37–39]. It is likely, therefore, that *MSH3* and *PMS2* modify XDP onset by altering the rate of somatic CCCTCT expansion.

We find greater levels of somatic expansion in all brain regions analyzed relative to levels in blood, supporting recent observations in two XDP patients [41]. Within the brain, we observe region-specific differences in the degree of repeat expansion that are reflected across the different XDP individuals, with cerebellum exhibiting the most stability and cortical structures tending to be the most unstable. Several other disease-associated microsatellite repeats are relatively stable in cerebellum [29,50,51]. Here, we show substantial correlation between brain region-specific levels of expansion of the XDP CCCTCT repeat and the *HTT* CAG repeat, as previously observed in a similar comparison between expansion of the *HTT* CAG repeat and of the *ATXN1* CAG repeat underlying spinocerebellar ataxia type 1 (SCA1) [29]. These data provide support for common proteins (*trans*-acting factors) that modify tissue-specific levels of somatic expansion of both the XDP and *HTT* repeats, as well as other disease-associated repeats. We also found that the SVA-associated CCCTCT repeat within the *LIPG* gene, not known to be associated with any disease, exhibited repeat length-dependent expansion that was low in cerebellum compared to other brain regions analyzed. Interestingly, the *LIPG* CCCTCT repeat exhibited less instability than the XDP CCCTCT repeat in most of the brain regions analyzed, despite its relatively longer repeat lengths, pointing to potential modification of CCCTCT repeat instability *in cis*. It is plausible that local chromatin structure at the *TAF1* SVA locus might predispose the CCCTCT repeat to expand, while at the *LIPG* SVA locus, expansion of the CCCTCT repeat is comparatively suppressed. In line with this idea, disease-associated short tandem repeats were found to be enriched at 3D chromatin boundaries; in contrast, matched non-disease-associated repeats did not exhibit such an enrichment [52]. Thus, insights into chromatin structural features at the XDP SVA locus relative to other non-disease associated SVAs may provide clues to the instability propensity of its CCCTCT repeat tract. We previously reported the high G-quadruplex-forming potential of the reverse orientation AGAGGG repeat in the XDP SVA sequence [4]; whether this plays a role in its repeat instability remains to be investigated. The *LIPG* gene is also expressed at low levels in brain tissues. Transcription has been proposed to play a role in promoting repeat instability [53], and therefore a low rate of transcription through the *LIPG* gene may contribute to the lower level of instability of the SVA-associated CCCTCT repeat within this gene.

Our analyses of repeat instability in different brain tissues do not immediately point to any clear correlation with brain regions implicated either through neuropathological or neuroimaging studies to be susceptible in XDP [10–16]. *e*.*g*. neuropathological changes have been described in tissues that include caudate, putamen, cortex and cerebellum [10], yet these regions encompass both the lowest (cerebellum) and highest (cortex) levels of expansion. However, the association of repeat instability and cellular vulnerability is currently challenging due to: 1) the lack of cell type-specific resolution of repeat instability; 2) limited XDP neuropathology data; 3) neurodegeneration, notably of MSNs [6–8]. In HD, GWAS studies have provided support for a two-step model of pathogenesis that depends both on the rate of somatic CAG expansion and repeat length threshold(s) needed to trigger a toxic process(es) [31]. Both the rate of repeat expansion and toxicity-eliciting threshold may differ by cell type, and as both instability and toxicity components are needed for pathogenesis, high levels of expansion do not necessarily predict cellular vulnerability, *e*.*g*. this provides a logical explanation for high levels of *HTT* CAG expansion seen in the liver yet the absence of obvious liver pathology [29]. A two-step model provides a framework for other repeat expansion diseases, and similarly can explain why the striatum is not the primary target of pathogenesis in SCA1 despite high levels of CAG expansion in that tissue [29]. We propose that this model can also be applied to XDP, predicting that somatic expansion of the CCCTCT repeat in certain cell types will elicit a toxic process(es) ultimately culminating in clinical disease. A full understanding of XDP pathogenesis will therefore entail dissecting both instability and toxicity components in specific cell types. Further, there is evidence for altered brain connectivity in XDP [12,14,15,54,55], providing added complexity such that repeat expansion in one cell-type may trigger functional deficits at the level of a neuronal circuit. Notably our results provide evidence for a landscape of somatic events that include both repeat expansions and contractions, highlighting the importance of cell type-specific level resolution to understand relationships with disease processes. Currently, the nature of the toxic species is unclear, with reduced TAF1 levels and novel TAF1 isoforms being plausible candidates [19,20,22–25]. The inverse correlation of *TAF1* mRNA levels with CCCTCT repeat length seen in blood [26] is consistent with a role of TAF1 levels in a pathological process triggered by CCCTCT repeat expansion. Finally, while the identification of the *MSH3* and *PMS2* genes as XDP onset modifiers provides strong support for repeat expansion as the upstream driver of a toxic process(es), TAF1 itself has been implicated in promoting genome integrity [56–58]. Therefore, it is possible that altered TAF1 function in the disease process may further impact the DNA repair processes that underlie repeat instability, *e*.*g*. our data hint at differences in the instability of a non-disease-associated repeat (*LIPG* CCCTCT tract) in some tissues. Thus, genome-wide analyses of DNA instability/integrity in XDP patient brain would be of interest.

The prediction from our data is that that somatic CCCTCT repeat expansion contributes to length-dependent clinical measures, such as AAO. Of note, in the current dataset we find no difference in somatic expansion measured in blood between patients reporting symptom onset as either being predominantly dystonia or parkinsonism (Additional File 2: Fig.S1). This is consistent with the similar relationship between repeat length and AAO in these two patient subsets (Additional File 2: Fig.S1). Larger sample numbers will be needed to provide sufficient power for further tests of associations of repeat instability with clinical endpoints such as AAO. In addition, further studies will be needed to understand the relationship between repeat instability in blood and brain that will inform tests of association of instability with clinical measures.

## Conclusions

These data demonstrate that the XDP CCCTCT repeat is unstable in somatic cells, exhibiting properties that are consistent with a role for somatic expansion in determining the timing of disease onset. Our data suggest further avenues of investigation aimed at understanding the dynamics of this repeat mutation and relationship to pathogenesis.

## List of abbreviations

XDP: X-linked dystonia-parkinsonism
SVA: SINE-VNTR-Alu
HD: Huntington’s disease
TAF1: TATA-binding-protein (TBP)-associated factor-1
TBP: TATA-binding-protein
LIPG: Lipase G, Endothelial Type
MSNs: Medium-spiny neurons
AAO: Age at Onset
AAD: age at death
GWAS: genome-wide association
IRBs: 
MGH: Massachusetts General Hospital
IRB: Institutional Review Board
gDNA: Genomic DNA
CCXDP: Collaborative Center for XDP
DIG: digoxygenin
BA9: frontal cortex Brodmann area 9
SbN: substantia nigra
ION: inferior olivary nucleus
RN: red nucleus
DCN: deep cerebellar nuclei
STh: subthalamic nucleus
SP-PCR: small pool-PCR
HTT: huntingtin
MSH3: MutS homolog
3: PMS2: PMS1 homolog2, Mistmatch repair system component
SCA1: Spinocerebellar ataxia type 1

## Declarations

### Ethics for approval and consent to participate

All participants provided written informed consent, and the study was approved by Massachusetts General Hospital (Boston, MA, USA) and Jose R. Reyes Memorial Medical Center (Manila, Philippines) Institutional Review Boards (IRBs). Post-mortem brain tissue from XDP patients was obtained in collaboration with the Collaborative Center for XDP (CCXDP), at Massachusetts General Hospital (Boston, MA, USA), Makati Medical Center (Makati City, Philippines), and the Sunshine Care Foundation (Panay, Philippines). All procedures related to the collection, processing, and use of XDP patient post-mortem brain tissues were approved by IRBs at Makati Medical Center (Makati City, Philippines) and Massachusetts General Hospital (Boston, MA, USA)

### Consent for publication

All authors consented to the publication of the manuscript.

### Availability of data and materials

The datasets used and/or analyzed during the current study are available from the corresponding authors on reasonable request. Requests for tissue specimens may be directed to.

### Authors’ contributions

Sample acquisition: EBP, MGM, CF-C, MSV-A, GPL, NGG-B, JBBL, PJA, TM-B, GA, MLS, JKD, CG, NS, ELM, MCA, CCED, DCB; Sample processing: EBP, MGM, LNC; Data acquisition: LNC, TG, AMM; Data analysis: LNC, AMM, RY, KC, TG, LJO, VCW; Data interpretation: LNC, AMM, LJO, VCW; Study conception: LJO, VCW; Design of the work: LNC, AMM, LJO, VCW; Manuscript drafting and revision: LNC, AMM, LJO, VCW.

## Acknowledgements

We thank Marcy MacDonald, Jim Gusella and Jong-Min Lee for helpful input and discussion. We also thank XDP patients and their families for their invaluable contributions to this research.

