## Additional File 1 for "Tissue-specific and repeat length-dependent somatic instability of the X-linked dystonia parkinsonism-associated CCCTCT repeat"

**Table S1: Primers and PCR Conditions**

| Repeat | Primer | Reagent | PCR conditions |
|---|---|---|---|
| XDP SVA Hexamer (Standard sizing) | 5’-[FAM]AGCAGTACAGTCCAGCTTTGGC-3′ | PrimeSTAR GXL | 94°C x 2 min; 30 x (98 °C x 10 s, 64°C x 35 s) |
| | 5′-CTCAAGCCTTATTACAATGCCAGT-3′ | | |
| *LIPG* SVA Hexamer (Standard sizing) | 5’-CAGCAGTACAGTCCAGCTTC -3 | PrimeSTAR GXL | 94°C x 2 min;30 x (98°C x 10 s, 60°C x 15 s, 69°C x 25 s) |
| | 5’-[FAM]AATAGAGTGTATCCGGTGAGC-3’ | | |
| XDP SVA Hexamer (Small pool-PCR) | 5’-[FAM]AGCAGTACAGTCCAGCTTTGGC-3′ | PrimeSTAR GXL | 94°C x 2 min; 35 x (98°C x10 s, 64°C x 35 s) |
| | 5′-CTCAAGCCTTATTACAATGCCAGT-3′ | | |

**Table S2: List of XDP patients with blood and postmortem brain tissue available**

| Patient | Blood Repeat Length | Brain Repeat Length |
|---|---|---|
| 17-012 | **42** | **43** |
| 17-019 | 42 | 42 |
| 18-002 | 41 | 41 |
| 18-003 | 42 | 42 |
| 18-004 | 41 | 41 |
| 19-009 | 39 | 39 |
| 19-012 | 52 | 52 |
| 19-013 | 41 | 41 |
| 19-015 | 36 | 36 |
| 19-017 | **44** | **45** |
| 19-018 | 45 | 45 |
| 19-019 | 38 | 38 |
| 19-020 | 43 | 43 |
| 19-021 | **37** | **36** |
| 20-024 | 46 | 46 |
| 20-025 | 38 | 38 |
| 20-026 | 39 | 39 |
| 21-027 | 43 | 43 |
| 21-028 | 37 | 37 |
| 21-029 | 42 | 42 |
| 21-031 | **47** | **48** |

**Table S3:** **XDP individuals with postmortem brain regions used for analyses of XDP CCCTCT repeat instability**

| Brain ID | BA9 | Cd | Cereb | Cin. Gyrus | DCN | Hip | Insula | ION | Lat. Th | Med. Th | Oc. Cx | Parietal Cx | Pu | RN | SbN | STh | TmP | *N* |
|---|---|---|---|---|---|---|---|---|---|---|---|---|---|---|---|---|---|---|
| 16-12 | X | X | X | X | X | X | X | | X | X | X | X | X | X | X | | X | 15 |
| 16-13 | X | X | X | X | X | X | X | X | X | X | X | X | X | X | X | X | | 16 |
| 17-01 | X | X | X | X | X | | X | | X | X | X | X | X | X | X | | X | 14 |
| 17-05 | X | X | X | X | X | X | X | X | X | X | X | X | X | X | X | X | X | 17 |
| 17-06 | X | X | X | X | X | X | X | X | X | X | X | X | X | X | X | | X | 16 |
| 17-10 | X | | X | X | X | X | X | | X | X | X | X | X | | | | X | 12 |
| 17-12 | X | X | X | X | X | X | X | X | X | X | X | X | X | X | X | | X | 17 |
| 17-13 | X | X | X | X | X | X | X | X | X | X | X | X | X | X | X | | X | 16 |
| 17-14 | X | | X | X | X | X | X | X | X | X | X | X | X | X | X | X | X | 16 |
| 17-17 | X | X | X | X | X | X | X | X | X | X | X | X | X | X | X | X | X | 17 |
| 17-19 | X | X | X | X | X | X | X | X | X | X | X | X | X | X | X | X | X | 18 |
| 18-001 | X | X | X | X | X | X | X | | X | X | X | X | X | | X | | X | 14 |
| 18-002 | X | | X | X | X | X | | | X | X | X | X | | | X | | X | 11 |
| 18-003 | X | X | X | X | X | X | X | | X | X | X | X | X | | X | | X | 14 |
| 18-004 | X | X | X | X | X | X | X | | X | X | X | X | X | | X | | X | 14 |
| 18-006 | X | X | X | X | X | X | X | | X | X | X | X | X | X | X | X | X | 16 |
| 19-007 | X | | X | | X | | | | | | X | | | | | | X | 5 |
| 19-008 | X | X | X | X | X | X | X | | X | X | X | X | X | | X | X | X | 15 |
| 19-009 | X | X | X | X | X | X | X | | X | X | X | X | X | | X | X | X | 16 |
| 19-010 | | | X | | | | | | | | | | | | | | | 1 |
| 19-011 | | | | | | | | | | | X | | | | | | | 1 |

| Brain ID | BA9 | Cd | Cereb | Cin. Gyrus | DCN | Hip | Insula | ION | Lat. Th | Med. Th | Oc. Cx | Parietal Cx | Pu | RN | SbN | STh | TmP | *N* |
|---|---|---|---|---|---|---|---|---|---|---|---|---|---|---|---|---|---|---|
| 19-012 | | | X | | | | | | | | | | | | | | | 1 |
| 19-013 | | | X | | | | | | | | | | | | | | | 1 |
| 19-014 | | | X | | | | | | | | | | | | | | | 1 |
| 19-015 | | | X | | | | | | | | | | | | | | | 1 |
| 19-016 | | | X | | | | | | | | | | | | | | | 1 |
| 19-017 | X | X | X | X | X | X | X | X | X | X | X | X | X | | X | | X | 16 |
| 19-018 | X | X | X | X | X | X | X | | X | X | X | X | X | | X | | X | 15 |
| 19-019 | | | X | | | | | | | | | | | | | | | 1 |
| 19-020 | | | X | | | | | | | | | | | | | | | 1 |
| 19-021 | | | X | | | | | | | | X | | | | | | | 2 |
| 19-022 | | | X | | | | | | | | | | | | | | | 1 |
| 20-024 | | | X | | | | | | | | | | | | | | | 1 |
| 20-025 | | | X | | | | | | | | | | | | | | | 1 |
| 20-026 | | | X | | | | | | | | | | | | | | | 1 |
| 21-027 | | | X | | | | | | | | | | | | | | | 1 |
| 21-028 | | | X | | | | | | | | | | | | | | | 1 |
| 21-029 | | | X | | | | | | | | | | | | | | | 1 |
| 21-030 | | | X | | | | | | | | | | | | | | | 1 |
| 21-031 | | | X | | | | | | | | X | | | | | | | 2 |
| 21-032 | | | X | | | | | | | | | | | | | | | 1 |
| **Total** | 21 | 17 | 40 | 20 | 21 | 19 | 19 | 9 | 20 | 20 | 24 | 20 | 19 | 11 | 19 | 8 | 20 | |

Brains tissues analyzed are: BA9 = frontal cortex Brodmann area 9, Cd=caudate, Cereb=cerebellum, Cin.Gyrus = cingulate gyrus, DCN = deep cerebellar nuclei, Hip = hippocampus, Insula = insular cortex, ION = inferior olivary nucleus, Lat.Th = lateral thalamus, Med.Th = medial thalamus, Oc. Cx = occipital cortex, Parietal Cx = parietal cortex, Pu = putamen, RN = red nucleus, SbN=substantia nigra, STh = subthalamic nucleus and TmP = temporal pole.

### Table S4: Pair-wise statistical comparisons of XDP CCCTCT expansion indices in blood and brain regions

| *P*-value | | | | | | | | | | | | | | | | | | |
|---|---|---|---|---|---|---|---|---|---|---|---|---|---|---|---|---|---|---|
| | **Blood** | **Cereb** | **Cd** | **SbN** | **ION** | **RN** | **Med.Th** | **Hip** | **Pu** | **Lat.Th** | **DCN** | **STh** | **Cin.Gyrus** | **BA9** | **Parietal Cx** | **Insula** | **TmP** | **Oc.Cx** |
| **Blood** | - | 2e-16 | 2.2e-09 | 1.6e-10 | 7.5e-05 | 4.5e-06 | 4.9e-11 | 1.6e-10 | 1.6e-10 | 4.5e-11 | 1.3e-11 | 2.8e-04 | 4.8e-11 | 1.3e-11 | 4.5e-11 | 1.5e-10 | 4.5e-11 | 4e-13 |
| **Cereb** | | - | ns | ns | ns | ns | ns | 0.05003 | ns | 1.3e-03 | 4e-03 | 0.03 | 1.7e-03 | 2.7e-05 | 1.5e-08 | 1.7e-08 | 2.6e-09 | 7.7e-10 |
| **Cd** | | | - | ns | ns | ns | ns | ns | ns | ns | ns | ns | ns | ns | 0.01 | 0.015 | 6e-03 | 1.9e-04 |
| **SbN** | | | | - | ns | ns | ns | ns | ns | ns | ns | ns | ns | ns | ns | 0.02 | 8.5e-03 | 3. 6e-04 |
| **ION** | | | | | - | ns | ns | ns | ns | ns | ns | ns | ns | ns | ns | ns | ns | ns |
| **RN** | | | | | | - | ns | ns | ns | ns | ns | ns | ns | ns | ns | ns | ns | 0.01 |
| **Med.Th** | | | | | | | - | ns | ns | ns | ns | ns | ns | ns | ns | ns | ns | 7e-03 |
| **Hip** | | | | | | | | - | ns | ns | ns | ns | ns | ns | ns | ns | ns | 0.01 |
| **Pu** | | | | | | | | | - | ns | ns | ns | ns | ns | ns | 0.04 | 8.5e-03 | 5.9e-04 |
| **Lat.Th** | | | | | | | | | | - | ns | ns | ns | ns | ns | ns | ns | 0.01 |
| **DCN** | | | | | | | | | | | - | ns | ns | ns | ns | ns | 0.04 | 1.7e-03 |
| **STh** | | | | | | | | | | | | - | ns | ns | ns | ns | ns | ns |
| **Cin.Gyrus** | | | | | | | | | | | | | - | ns | ns | ns | ns | ns |
| **BA9** | | | | | | | | | | | | | | - | ns | ns | ns | ns |
| **Parietal Cx** | | | | | | | | | | | | | | | - | ns | ns | ns |
| **Insula** | | | | | | | | | | | | | | | | - | ns | ns |
| **TmP** | | | | | | | | | | | | | | | | | - | ns |
| **Oc.Cx** | | | | | | | | | | | | | | | | | | - |

Table shows the results of pairwise Wilcoxon rank sum tests comparing expansion indices in each tissue with Bonferroni-corrected P values. Note that the comparisons are not equally powered as the number of samples for each tissue differs (Table S3).

ns = not significant.

**Table S5: Frequencies of repeat length changes relative to the modal allele from single molecule SP-PCR data**

| Repeat length change | Frequency of events (%) | | | | | | | | |
|---|---|---|---|---|---|---|---|---|---|
| | 17-17 | | | | 19-008 | | | | 18-006 |
| | Cd | Pu | Cereb | Oc. Cx | Cd | Pu | Cereb | Oc. Cx | Oc. Cx |
| ≥-35 | 0.41 | 0 | 0.82 | 0 | 0 | 0 | 0 | 0 | 0 |
| ≥-30 | 1.64 | 0 | 0.82 | 0.47 | 0 | 0 | 0 | 0 | 0 |
| ≥-25 | 2.46 | 0 | 0.82 | 1.89 | 0 | 0 | 0 | 0 | 0 |
| ≥-20 | 2.46 | 0 | 1.65 | 2.84 | 0 | 0 | 0 | 0.59 | 0 |
| ≥-15 | 2.88 | 0 | 2.47 | 3.79 | 0 | 0 | 0.7 | 2.38 | 0 |
| ≥-10 | 2.88 | 0 | 7.43 | 5.68 | 0.7 | 0 | 0.7 | 2.97 | 0 |
| ≥-5 | 4.11 | 0 | 9.92 | 10.9 | 0.7 | 0.6 | 0.7 | 2.97 | 0.54 |
| ≥-1 (all contractions) | 29.21 | 22.4 | 45.45 | 32.23 | 16.9 | 23.7 | 16.31 | 26.19 | 17.58 |
| 0 (modal allele) | 28.39 | 34.69 | 24.79 | 15.63 | 28.87 | 26.83 | 28.37 | 20.24 | 24.17 |
| ≥+1 (all expansions) | 42.38 | 42.85 | 29.75 | 52.13 | 54.22 | 49.39 | 55.31 | 53.57 | 58.24 |
| ≥+5 | 2.46 | 3.4 | 14.04 | 18.48 | 7.74 | 6.7 | 4.96 | 9.52 | 14.28 |
| ≥+10 | 1.64 | 1.36 | 14.04 | 8.05 | 0.7 | 2.43 | 0 | 2.97 | 5.49 |
| ≥+15 | 1.23 | 1.36 | 13.22 | 5.68 | 0 | 2.43 | 0 | 0 | 2.19 |
| ≥+20 | 0.41 | 0 | 12.39 | 3.31 | 0 | 2.43 | 0 | 0 | 1.09 |
| ≥+25 | 0.41 | 0 | 12.39 | 2.36 | 0 | 2.43 | 0 | 0 | 0.54 |
| ≥+30 | 0.41 | 0 | 12.39 | 2.36 | 0 | 2.43 | 0 | 0 | 0 |
| ≥+35 | 0 | 0 | 12.39 | 0.94 | 0 | 2.43 | 0 | 0 | 0 |
| ≥+40 | 0 | 0 | 12.39 | 0.94 | 0 | 2.43 | 0 | 0 | 0 |
| ≥+45 | 0 | 0 | 0.82 | 0.47 | 0 | 2.43 | 0 | 0 | 0 |

Frequencies of repeat length changes relative to the modal allele meeting various thresholds. Negative repeat length changes (contractions) of at least one repeat (≥-1) represent all contraction events, with progressively lower frequencies of contractions of ≥5, ≥10, ≥15, ≥20, ≥25, ≥30 and ≥35 repeats. Positive repeat length changes (expansions) of at least one repeat (≥+1) represent all expansion events, with progressively lower frequencies of expansions of ≥5, ≥10, ≥15, ≥20, ≥25, ≥30, ≥35, ≥40 and ≥45 repeats.

**Table S6: XDP individuals with postmortem brain regions used for analyses of *LIPG* CCCTCT repeat instability**

| **Brain ID** | **Cereb** | **Cd** | **Hip** | **BA9** | **TmP** | **Oc. Cx** | ***LIPG* repeat lengths analyzed [a]** |
|---|---|---|---|---|---|---|---|
| **16-12** | X | X | X | X | X | X | 48,56 |
| **16-13** | X | | | | | | 44 |
| **17-01** | X | | | | | | 49,57 |
| **17-05** | X | | | | | | 55 |
| **17-06** | X | | | | | | 61 |
| **17-10** | | | | X | | | 47 |
| **17-13** | X | X | X | X | X | X | 45,57 |
| **17-19** | X | X | X | X | X | X | 49,60 |
| **18-001** | X | | | | | | 39 |
| **18-002** | X | | X | X | X | X | 40,67 |
| **18-003** | X | | | | | | 58 |
| **18-004** | X | X | X | X | X | X | 53,64 |
| **19-008** | X | | | | | | 50 |
| **19-009** | X | X | X | X | X | | 49,57 |
| **19-010** | X | | | | | | 56 |
| **19-012** | X | | | | | | 53,60 |
| **19-013** | X | | | | | | 52 |
| **19-014** | X | | | | | | 69 |
| **19-017** | X | | | | | | 53 |
| **19-018** | X | | | | | | 60 |

| | | | | | | | |
|---|---|---|---|---|---|---|---|
| **19-019** | X | | | | | | 63 |
| **19-020** | X | | | | | | 52 |
| **19-021** | X | | | | | | 51,61 |
| **19-022** | X | | | | | | 50,55 |
| ***N*** | 23 | 5 | 6 | 7 | 6 | 5 | |

[a] only one allele is indicated either because only one major allele was detected, indicating homozygosity or because the proximity of the two alleles precluded accurate analysis of expansion peaks of the shorter allele.

**Table S7: Pair-wise statistical comparisons of *LIPG* CCCTCT expansion indices in brain regions**

| Brain region | BA9 | Cd | Cereb | TmP | Hip | Oc. Cx |
|---|---|---|---|---|---|---|
| **BA9** | - | ns | 2.8e-08 | ns | ns | ns |
| **Cd** | | - | 1.2e-08 | ns | ns | ns |
| **Cereb** | | | - | 6.5e-09 | 9.7e-09 | 1.3e-07 |
| **TmP** | | | | - | ns | ns |
| **Hip** | | | | | - | ns |
| **Oc. Cx** | | | | | | - |

Table shows the results of pairwise Wilcoxon rank sum tests comparing expansion indices in each tissue with Bonferroni-corrected P values.

ns = not significant.
