## Additional File 2 for "Tissue-specific and repeat length-dependent somatic instability of the X-linked dystonia parkinsonism-associated CCCTCT repeat"

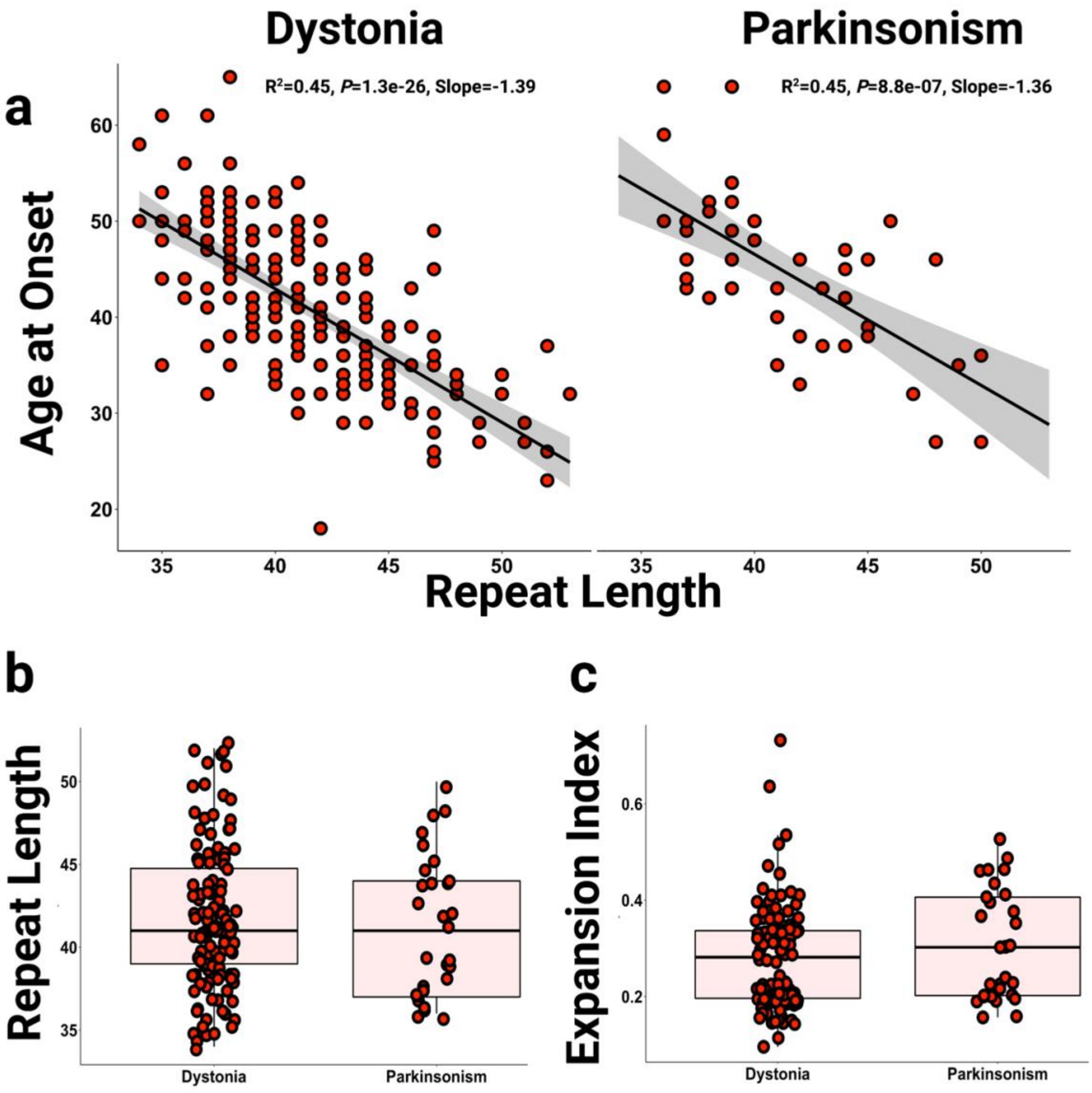


**Fig S1. Repeat-related outcomes in individuals exhibiting dystonia or Parkinson's symptoms at onset. a** Inverse correlations between CCCTCT repeat length (in blood) with AAO in individuals expressing dystonia (n=194) or parkinsonism (n=43) at onset. Grey shaded areas show 95% confidence interval**. b** Distribution of repeat length (in blood) in individuals expressing dystonia (n=194) or parkinsonism (n=43). **c** Distribution of expansion indices (in blood) in individuals expressing dystonia (n=122) or parkinsonism (n=29). Box-whisker plots show median ± interquartile range (IQR) and red dots show values in individual patient samples. **b-c** showed no statistical differences between groups.

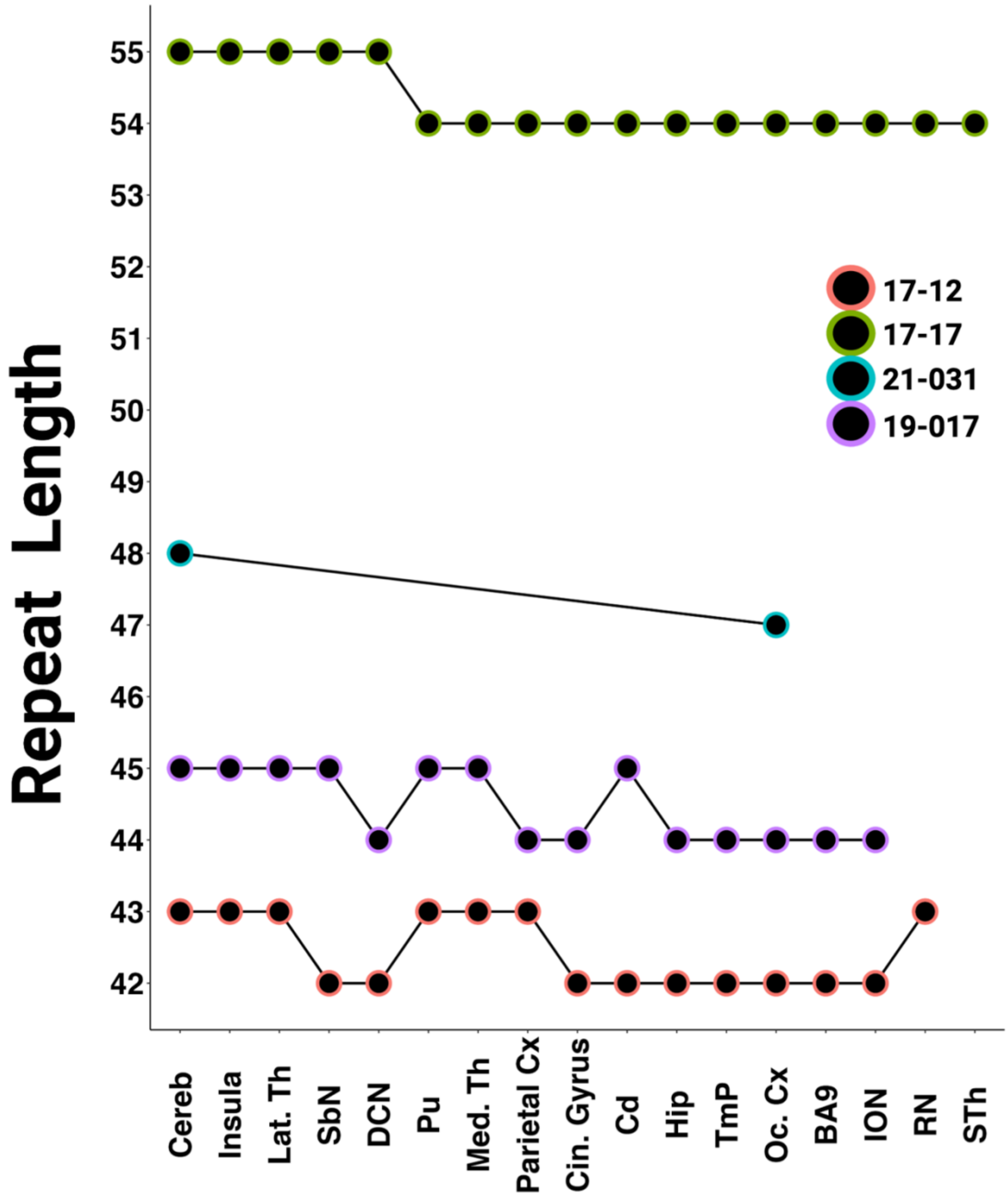


**Fig S2: Modal repeat length variation in brain regions**. Of the 23 postmortem samples in which multiple brain regions were analyzed, 4 (17-012, 17-17, 19-017 and 21-031) exhibited variation by one repeat unit in the modal repeat length.

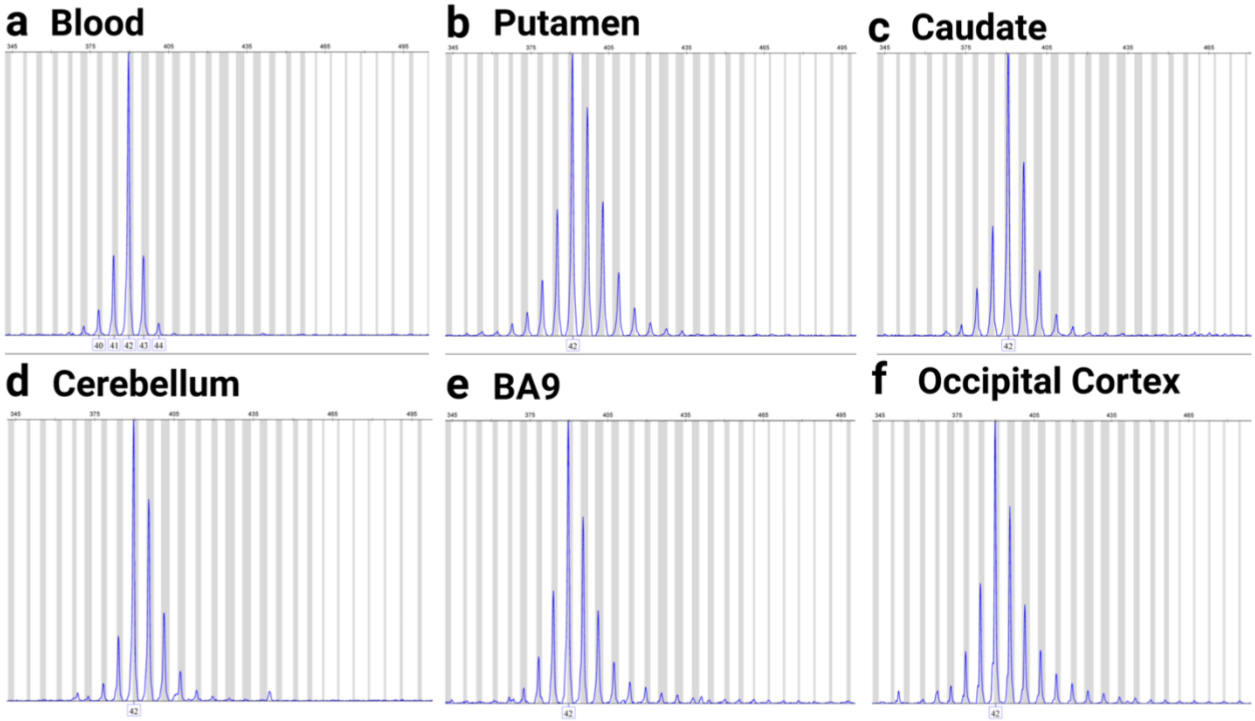


**Fig S3: Examples of XDP CCCTCT repeat GeneMapper traces from different tissues. a-f** GeneMapper traces of blood and different brain tissues from individual 17-19. "x" axis shows base pair size and peak height respectively. Indicated on the "x"-axes are assigned repeat lengths based on standard repeat controls.

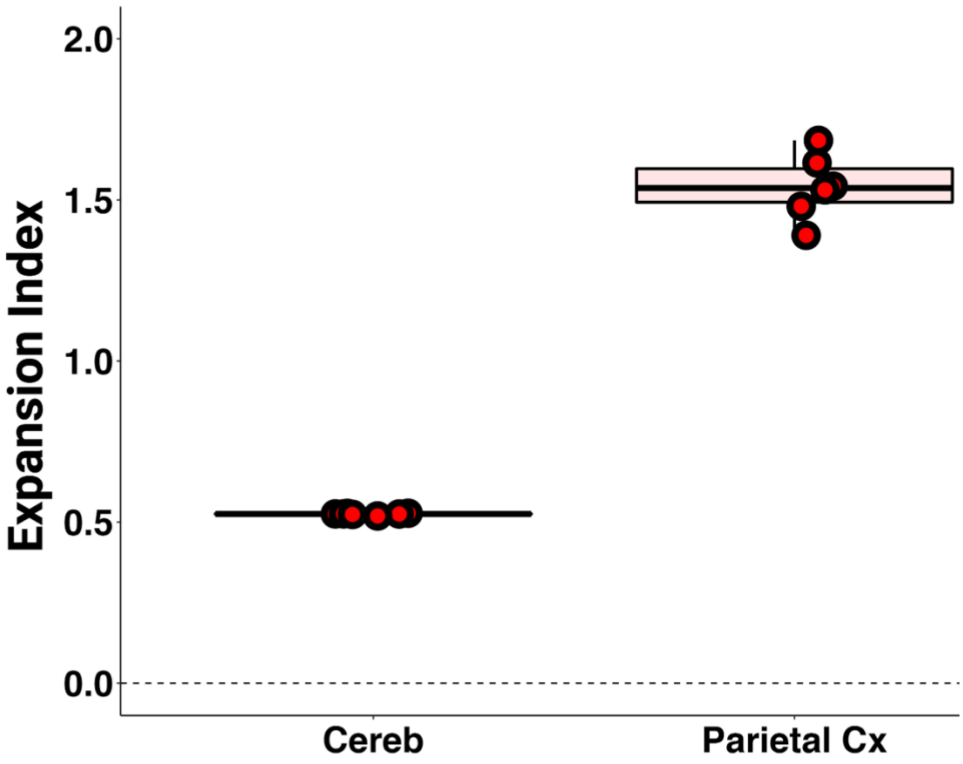


**Fig. S4: Technical variability in expansion index values.** Expansion indices were derived from 5 independent XDP CCCTCT repeat PCRs from samples 19-022 (Cereb) and 18-001 (Parietal Cx).

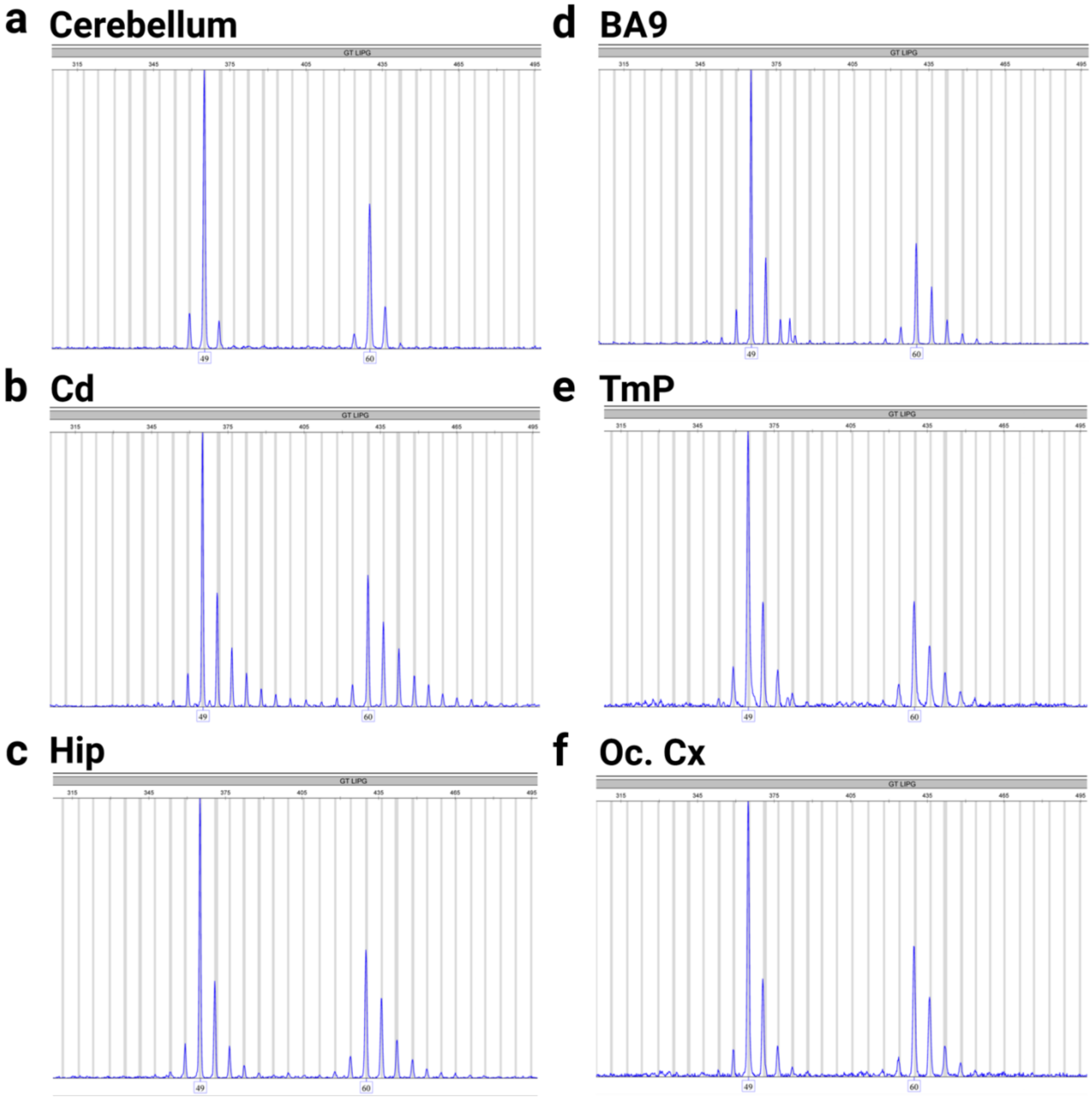


**Fig S5: Examples of *LIPG* CCCTCT GeneMapper traces from different tissues.**

**a-f** GeneMapper traces of different brain tissues from individual 17-19. “x” axis shows base pair size and peak height respectively. Indicated on the “x”-axes are assigned repeat lengths based on base pair size.
